# Evaluating the diagnostic capabilities of nanopore sequencing for *Borrelia burgdorferi* detection in blacklegged ticks

**DOI:** 10.1101/2025.08.26.672273

**Authors:** Jacob Cassens, Evan J. Kipp, Lexi E. Frank, Peter A. Larsen, Jonathan D. Oliver

## Abstract

Ticks pose substantial threats to public health. Blacklegged ticks (*Ixodes scapularis*) are responsible for most tick-borne diseases in the US, transmitting seven human pathogens. Molecular surveillance for tick-borne pathogens has been outpaced by their emergence, revealing a critical need to develop agnostic strategies that characterize emerging and putative pathogens. Oxford Nanopore Technology’s nanopore adaptive sampling (NAS), an approach that selectively enriches or depletes for target genomes or genetic loci, provides an opportunity to generate real-time genomic insights into tick-borne pathogens. In the current study, we performed PCR and NAS on pooled *Borrelia burgdorferi-*infected and -uninfected ticks to evaluate the diagnostic capability of NAS. We found that NAS generates extensive datasets on tick-borne pathogens from individual ticks that aid in distinguishing true and false positive samples. Using a pooled approach consisting of whole genomic DNA from 168 total ticks multiplexed over seven sequencing experiments, our results indicated that NAS is extremely specific (0.97 [95% CI: 0.93, 1.00]) with moderate sensitivity (0.48 [95% CI: 0.41, 0.55]), suggesting a strong capacity to confirm *B. burgdorferi* when present at the expense of an elevated false-negative rate. We found that quality-based filtering of sequence data has a profound influence on diagnostic metrics, emphasizing the need to optimize pooling strategy, wet-lab procedures, and bioinformatic pipelines to enhance the sensitivity of NAS for detecting tick-borne pathogens.

**Author summary:** Ticks pose substantial threats to public health. In the United States, the blacklegged tick (*Ixodes scapularis*) can transmit seven (known) human pathogens and is responsible for most tick-borne disease cases. The most prevalent tick-borne pathogen, *Borrelia burgdorferi*, causes Lyme disease, which is consequently the most common tick-borne disease in the United States. As blacklegged ticks continue to expand across the United States, it is imperative to develop rapid molecular surveillance tools that agnostically detect emerging and putative pathogens. In the current study, we employ Oxford Nanopore Technology’s nanopore adaptive sampling on PCR-infected and PCR-uninfected blacklegged ticks to determine the diagnostic capability of this technology for rapid *Borrelia burgdorferi* detection. Our results suggest that nanopore adaptive sampling can generate extensive sequence datasets on user-specified target reference genomes. We demonstrate that nanopore adaptive sampling is extremely specific, capable of confirming *B. burgdorferi* presence when detected, yet lacks sensitivity, leading to a high false negative rate. We highlight the methodological limitations that undoubtedly led to this lower sensitivity and provide future research directions for enhancing nanopore adaptive sampling as a real-time, unbiased molecular tool for tick-borne pathogen surveillance.

## Introduction

Blacklegged ticks (*Ixodes scapularis*) are the primary source of locally acquired vector-borne disease in the US [1]. These ticks are known to transmit at least seven known human pathogens, spanning bacterial, viral, and protozoan taxa [2]. *Borrelia burgdorferi*, the etiological agent of Lyme disease, causes over 40,000 reported cases annually, which is likely a ten-fold underestimate of the true burden [3]. Lyme disease incidence mirrors the geographic distribution of blacklegged ticks, with primary endemic foci in the Mid-Atlantic, Northeast, and Midwest regions of the US [4]. Treatment of tick-borne disease in the US causes over $1.3 billion in direct medical costs annually, not including social costs or quality-of-life impacts [5]. These factors, together with the lack of effective control methods, have created the impetus to improve our surveillance systems for effective mitigation and prevention of tick-borne disease, particularly given the complex ecology of tick-pathogen interactions.

Blacklegged ticks are obligate blood-feeding ectoparasites, requiring a blood meal at each life stage for development (immatures) and reproduction (females). These ticks will quest on vegetation to seek animal hosts, attach, and feed until repletion. This feeding pattern enables the horizontal maintenance of tick-borne pathogens (TBPs), where transmission can occur through feeding on previously infected animals or co-feeding in proximity to infected ticks [6,7]. Blacklegged ticks have been expanding throughout the US over the past two decades, likely owing to shifts in animal hosts and increased habitat suitability [8–11]. In the wake of their expansion, TBPs have experienced similar shifts in presence and distribution, coincident with the steady rise in tick-borne disease incidence over the same period [12–14]. However, our current strategies to monitor shifts in TBP distribution and characterize their genetic diversity in wild ticks remain limited. Conventional TBP detection relies on amplification techniques targeting a single pathogen, with contemporary multiplex amplification approaches enabling the detection of multiple pathogens simultaneously [15,16]. Ticks are commonly infected with numerous microorganisms, rendering targeted amplification assays limited in their ability to characterize the diversity of microbes present in ticks agnostically [17,18]. As such, novel approaches that leverage next-generation sequencing, while preserving targeted TBP capture, represent an opportunity to enhance current surveillance approaches.

Oxford Nanopore Technologies (ONT) nanopore sequencing is a third-generation sequencing technology that uses biological nanopores distributed across a flow cell to sequence native genomic DNA (Oxford Nanopore Technologies, Oxford, United Kingdom). A constant ionic current is sent across the flow cell, where individual nucleotides sent through a nanopore cause distinct changes in electrical current that enable single-molecule real-time sequencing [19]. Nanopore adaptive sampling (NAS) couples nanopore sequencing with the real-time bioinformatic enrichment or depletion of target sequences, specified by the user in an indexed reference file [20]. Briefly, multiple reference genomes or target sequences are concatenated into the indexed reference file. As DNA fragments are sequenced, the initial bases of each read are aligned to the reference file for enrichment or depletion. If enrichment is chosen, the sequenced bases are aligned to the reference file using minimap2 [21], and the system uses the resulting alignment score to determine whether the strand is aligned inside or outside the target regions. A sufficient match results in the DNA fragment under question continuing to be sequenced. Conversely, if there is an insufficient match, the individual nanopore responsible for sequencing that fragment will reverse polarity and eject the molecule, freeing the nanopore for processing a new fragment. In this way, NAS serves as a dynamic platform to generate robust sequence data in metagenomic or targeted surveillance approaches. Previous research has demonstrated the utility of NAS for detecting host blood meals in insects [22], characterizing microorganisms in ticks [23] and insects [24], identifying host species from fecal matter [25], and rapidly monitoring infectious disease outbreaks [26].

In the current study, we evaluate the diagnostic capability of NAS for *B. burgdorferi* detection in naturally infected blacklegged ticks. Our objective was to compare diagnostic performance to conventional amplification techniques, assess the technical feasibility of NAS, and elucidate the influence of bioinformatic decisions on diagnostic metric ascertainment (Figure 1). We show that pooled NAS libraries can generate extensive sequence datasets on *B. burgdorferi*, with extremely high specificity and positive predictive value (PPV). We discuss our findings in light of the trade-offs between diagnostic metrics for TBPs and provide recommendations for future research to expand on our experimental approach. This study provides a foundation for future surveillance strategies that bridge traditional amplification techniques with genomic sequencing to detect known, emerging, and putative TBPs in real-time.

**Figure 1.**
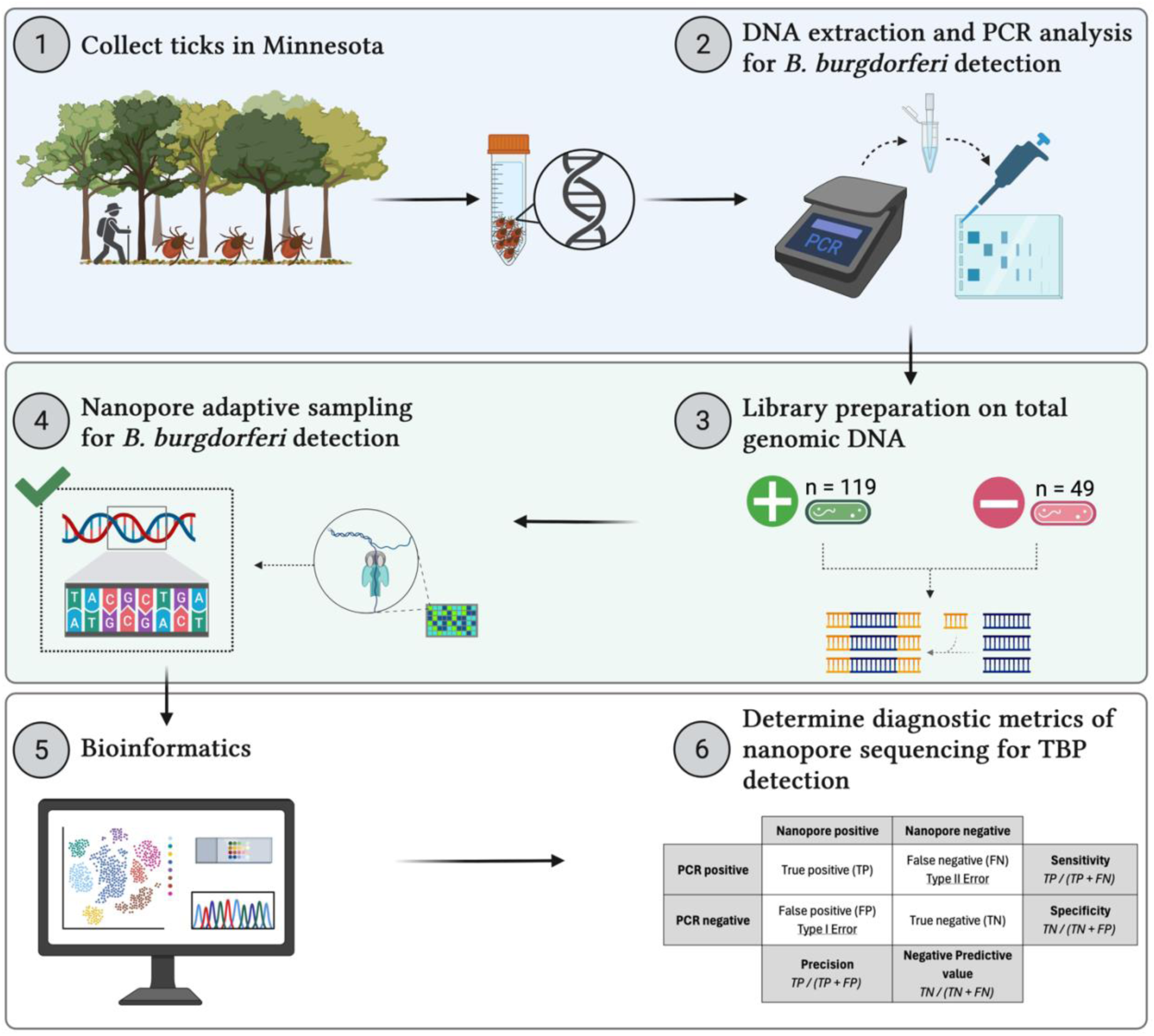
Diagram displaying the experimental workflow. (1) Questing blacklegged ticks were collected across three counties in Minnesota using 1 m2 drag cloths and placed in 70% EtOH for transportation to the laboratory. (2) Ticks were surface sterilized, morphologically identified, and total genomic DNA extracted. Nested PCR analysis for *B. burgdorferi* was performed on each extract. (3) nPCR-positive and negative ticks were randomly selected, and their genomic DNA was used in library preparation to generate seven individual libraries containing 24 barcoded and pooled ticks for nanopore sequencing. (4) Nanopore adaptive sampling (NAS) files were created to enrich for all known bacterial pathogens vectored by blacklegged ticks. Sequencing was performed on seven individual flow cells for 72 hours. (5) POD5 files were rebasecalled, and the passed FASTQ files were used for downstream bioinformatic analysis. (6) Putative NAS infection status was coupled with their PCR status to determine the diagnostic capability of NAS for *B. burgdorferi* detection in blacklegged ticks.

## Results

### Tick Collection and PCR Analysis

Over the course of May to July in 2023 and 2024, more than 800 ticks were collected across 115 surveyed linear kilometers at three sites in Minnesota (Table 1). 275 were morphologically identified as adult blacklegged ticks (*Ixodes scapularis*) and subjected to molecular screening for *B. burgdorferi*. Among those, 172 (62.5%) were positive and 103 (37.5%) were negative, confirmed in duplicate nested PCR (nPCR) experiments to validate the unexpectedly high prevalence. DNA concentrations ranged from 2.1 ng/μL to 112 ng/μL before vacuum centrifuge concentration, which concentrated all DNA extracts to >50 ng/μL.

**Table 1.**
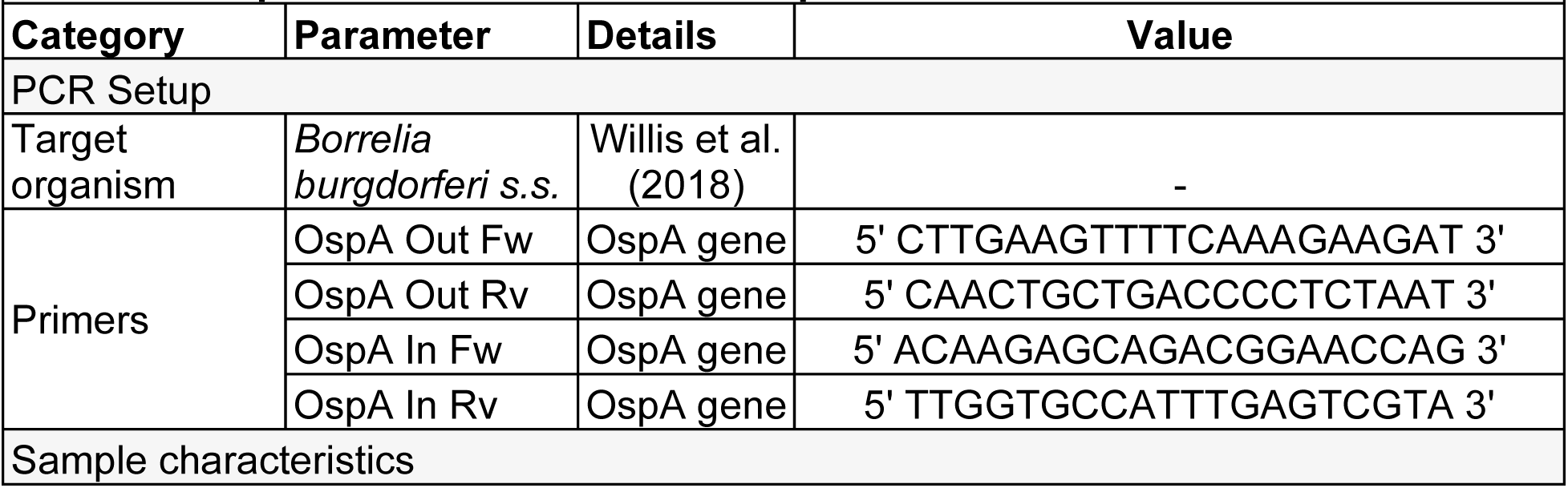
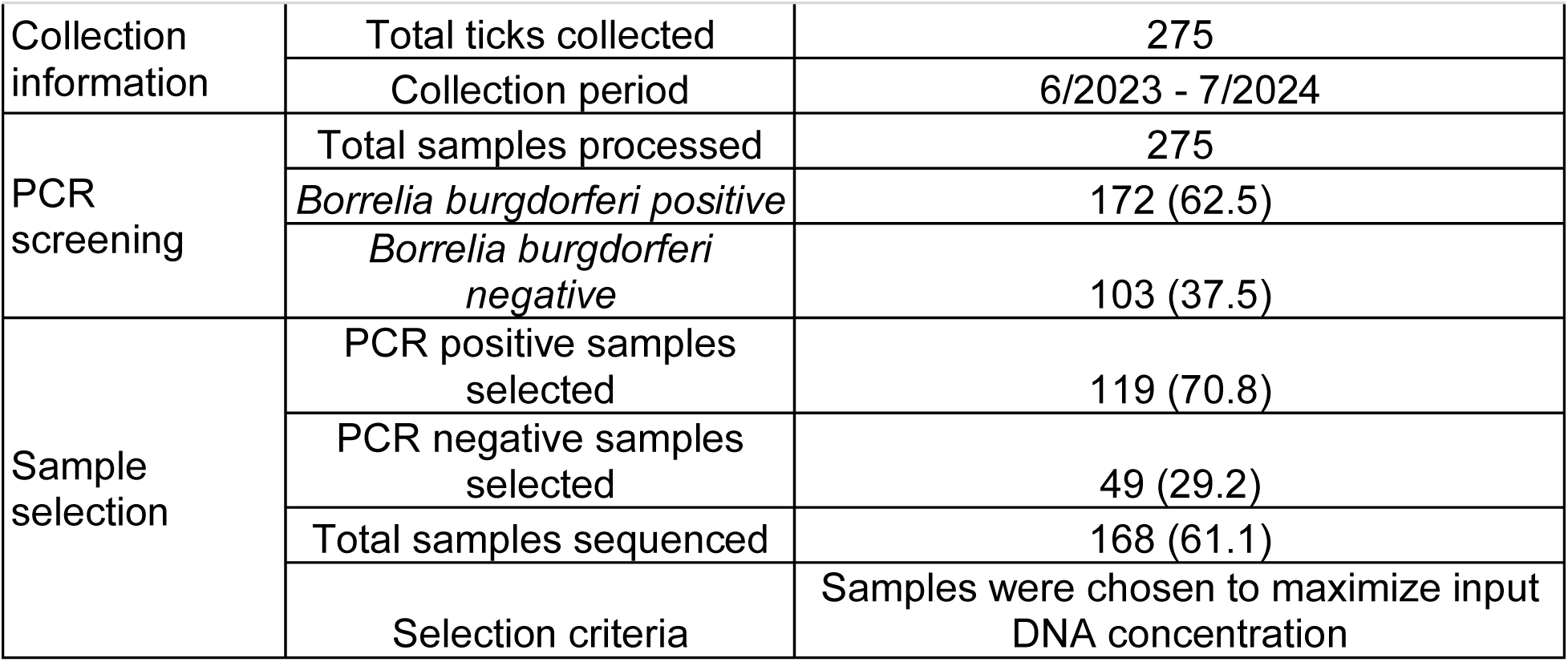
Sample characteristics and PCR primer details.

### Nanopore Sequencing Performance

The number of pores available for sequencing at the beginning of each run ranged from 850 to 1600 per flow cell, with no discernible effect on total data yield (Table 2). Across the seven sequencing experiments (1.98 μg to 3.01 μg total tick gDNA pooled per experiment), a total of 266 million (M) reads and 103 Gb were generated, averaging 38 M reads and 14 Gb per experiment (Table 2). Sequence experiment yields varied widely from 25 M to 57 M reads and 11 to 21 Gb (Table 2). The variation in sequencing yield was not correlated with the initial number of available pores on flow cells, suggesting that DNA quality, concentration, and total input library concentration likely hold primary influence. Mean read averaged 392 bp with a Q score of 8.5 and an N50 of 438 bp (Table 2). This likely reflects NAS’s rejection of many non-target sequences that yield lower average read lengths and N50 scores compared to conventional non-targeted nanopore sequencing. Sequence summary text files from each experiment enabled the calculation of adaptive sampling efficiency. In total, only 0.1% of sequenced bases were on-target (mapped to the *B. burgdorferi* reference genome during NAS), indicating that the majority of sequenced reads and bases were either off-target or mapped to other TBP genomes in the enrichment file. Generally, data output from each sequencing experiment plateaued as run time increased, with the number of active pores sharply declining after 50 hours of sequencing (Figure 2). Read length and quality remained constant throughout the sequencing experiments, while the number of reads per channel on flow cells displayed uniform distribution across the 512 channels, with a few channels showing either no activity or extremely high activity (Figure 2). Sequence experiment characteristics for each run are available in Supplementary File S1. FASTQ files from each sequencing experiment were partitioned into nPCR-infected classes to compare sequence yield and characteristics across classes and filtering criteria. Infected ticks generated more sequences and therefore greater absolute numbers of bases, with largely invariant quality, N50, and average length statistics in comparison to data from uninfected control ticks (Table 3). Infected ticks produced longer maximum read lengths, and increasing the quality filtering stringency reduced the number of sequences by 82% (from raw to Q10) and 62% (from Q10 to Q20), respectively, across both infected and control ticks (Table 3).

**Figure 2.**
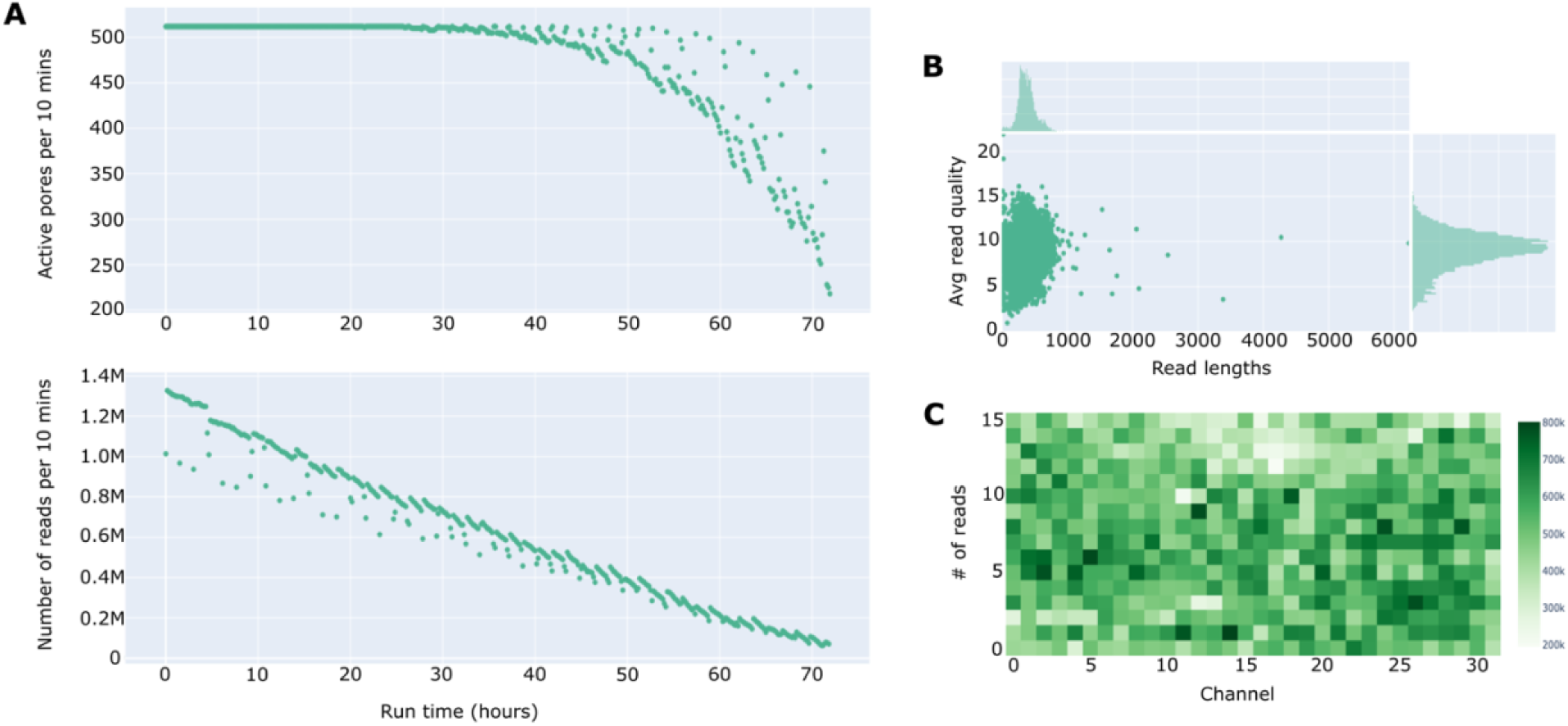
Sequence summary statistics distributions for all nanopore adaptive sampling experiments generated with NanoPlot. The number of active pores and the number of reads generated across all experiments over the 72-hour sequencing period are shown in (A). The relationship between average read length and quality, with their respective distributions, is shown in (B). The number of reads generated per channel, across the 512 channels distributed across a flow cell, is shown in (C).

**Table 2.** General nanopore experiment statistics generated using sequence summary text files with NanoStat and python. Sequencing experiments were run for 72 hours. Each sequencing experiment included 24 tick genomic DNA extracts, barcoded and pooled for sequencing using the SQK-NBD114.24 kit.

|  |  |  |  |  |  |  |  |  |  |
| --- | --- | --- | --- | --- | --- | --- | --- | --- | --- |
| **Experiment** | **Available pores** | **# of reads (M)** | **# of bases (Gb)** | **N50 (bp)** | **Mean read length (bp)** | **Mean read quality** | **On-target reads\*** | **On-target bases\* (bp)** | **% bases on target** |
| Run 1 | 1519 | 25.69 | 11.46 | 522 | 446.2 | 8.8 | 1,326 | 1,044,469 | 0.01% |
| Run 2 | 1543 | 38.79 | 17.21 | 514 | 443.6 | 9.0 | 1,724 | 1,322,073 | 0.01% |
| Run 3 | 1370 | 32.30 | 14.51 | 523 | 449.2 | 8.7 | 1,154 | 974,847 | 0.01% |
| Run 4 | 1525 | 57.97 | 21.03 | 387 | 362.8 | 8.4 | 2,714 | 2,293,490 | 0.01% |
| Run 5 | 850 | 31.31 | 11.27 | 402 | 360.1 | 8.0 | 433 | 534,729 | 0.001% |
| Run 6 | 1249 | 47.56 | 16.51 | 347 | 347.2 | 8.4 | 2,086 | 729,514 | 0.01% |
| Run 7 | 1040 | 33.30 | 11.27 | 371 | 338.4 | 8.0 | 1,629 | 521,231 | 0.01% |
| **Avg.** | 1300 | 38.13 | 14.75 | 438 | 392.5 | 8.5 | 1,581 | 1,060,050 | 0.01% |
| **Total** | - | 266.9 | 103.3 | 414 | 386.9 | 8.4 | 11,066 | 7,420,353 | 0.06% |

\*: On-target reads and bases represent those mapping to the *B. burgdorferi* reference genome during adaptive sampling. Off-target reads and bases are those mapping to other reference genomes included in enrichment file.

**Table 3.** Comparison of raw and filtered FASTQ file summary statistics between PCR positive and negative samples. Values represent averages for each nPCR status group.

| nPCR status | n | # of seqs (K) | Sum length (Mb) | Avg length (bp) | Max length (kb) | N50 (bp) | N50 num | Q20(%) | Avg quality | GC(%) |
| --- | --- | --- | --- | --- | --- | --- | --- | --- | --- | --- |
| **Raw** |  |  |  |  |  |  |  |  |  |  |
| Control | 49 | 493.2 | 114.9 | 230.4 | 13.3 | 250.3 | 1049.7 | 81.3 | 16.2 | 45.4 |
| Infected | 119 | 590.4 | 136.7 | 230.2 | 14.9 | 246.8 | 1074.5 | 81.6 | 16.4 | 45.6 |
| **Q10 & >300 bp** |  |  |  |  |  |  |  |  |  |  |
| Control | 49 | 95.0 | 37.9 | 382.4 | 13.3 | 366.6 | 925.4 | 82.3 | 16.7 | 45.4 |
| Infected | 119 | 109.1 | 42.8 | 379.7 | 14.8 | 361.9 | 952.8 | 82.7 | 16.9 | 45.5 |
| **Q20 & >300 bp** |  |  |  |  |  |  |  |  |  |  |
| Control | 49 | 35.6 | 14.1 | 377.8 | 8.6 | 362.2 | 560.6 | 93.0 | 22.7 | 44.5 |
| Infected | 119 | 41.4 | 16.2 | 373.2 | 10.3 | 355.2 | 588.0 | 92.2 | 22.6 | 44.0 |

### Sequence Mapping and Filtering Analysis

Raw and filtered FASTQ files were mapped to the *B. burgdorferi* reference genome (B31; GCF_000008685) to ascertain the infection status of individual blacklegged ticks using NAS. Individual ticks were deemed NAS positive by the number of reads mapping to the reference genome, initially using any reads as putative positivity and subsequently removing any putative NAS-positive ticks if they did not contain more than 10 mapped reads. Raw sequencing files revealed an infection prevalence of 45.8% among all ticks, demonstrably lower than the nPCR infection prevalence of 70%. Filtering FASTQ files for higher quality scores and longer read lengths led to decreased putative infectivity, dropping to 40.4% for Q10 and 36.9% for Q20, respectively. Additional filtering for more than 10 mapped reads led to further reductions in putative infection detection to 33.3%, 28.6%, and 25.6% across the three quality categories, respectively. A detailed workflow of the bioinformatic analysis is displayed in Figure 3.

**Figure 3.**
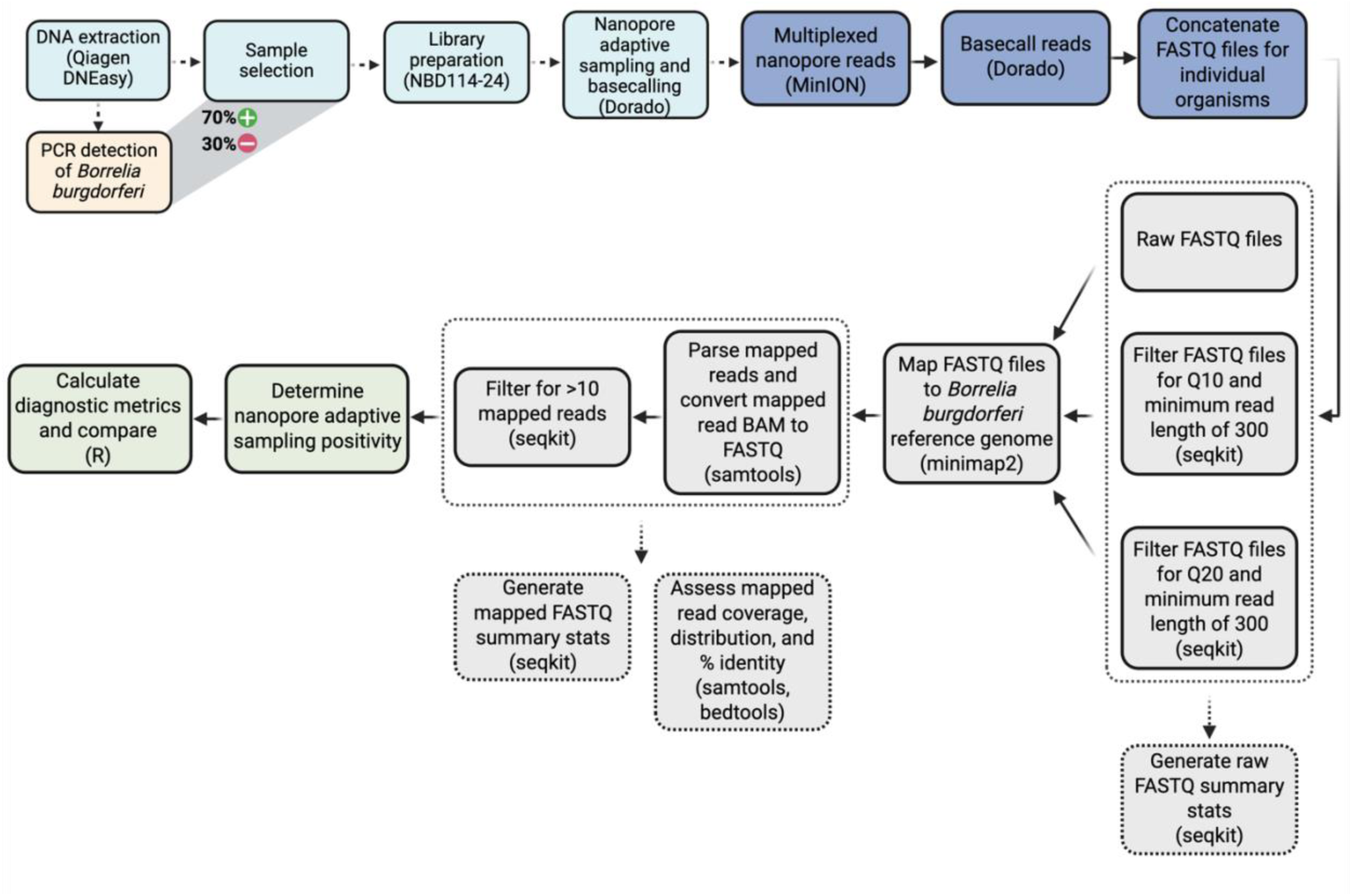
Diagram demonstrating the bioinformatic workflow and pipeline used to analyze NAS data. Parentheses reference the wet-lab kits used at each step and/or bioinformatic tools for analyzing sequence data.

Following the determination of putative NAS infection status, FASTQ files were partitioned according to their putative NAS and nPCR status for comparison. Putative true positive samples (NAS+ and nPCR+) outnumbered putative false positive samples (NAS+ and nPCR-) across all filtering thresholds (Table 4). Filtering for higher quality scores and longer read lengths resulted in successive reductions in putative false positives, and filtering FASTQ files for more than 10 mapped reads eliminated all putative false positives (Table 4). Across all filtering categories, putative true positive sequence characteristics differed substantially from putative false positives, demonstrating a greater number of sequences, bases, read lengths, and N50 lengths, despite nearly identical quality scores (Table 4). Mapped reads from putative true positive samples displayed much lower GC percentages (∼28%), often an order of magnitude lower, consistent with the *B. burgdorferi* genome’s average GC content (28%). Mapped reads were separated according to their mapped location to ascertain the number of reads and coverage across the chromosomal DNA and extrachromosomal plasmid DNA (Figure 4). Mapped reads were unevenly distributed across the genome, with the majority of reads mapping to the chromosomal DNA scaffold (Figure 5). Filtering for higher quality and longer mapped read length reduced the total number and coverage of chromosomal and plasmid reads, whereas removing putative positives with less than 10 mapped reads increased the number and coverage of chromosomal and plasmid reads. Putative true positive mapped reads demonstrated greater percent identity, mean read length, and coverage across the *B. burgdorferi* genome (Figure 5).

**Figure 4.**
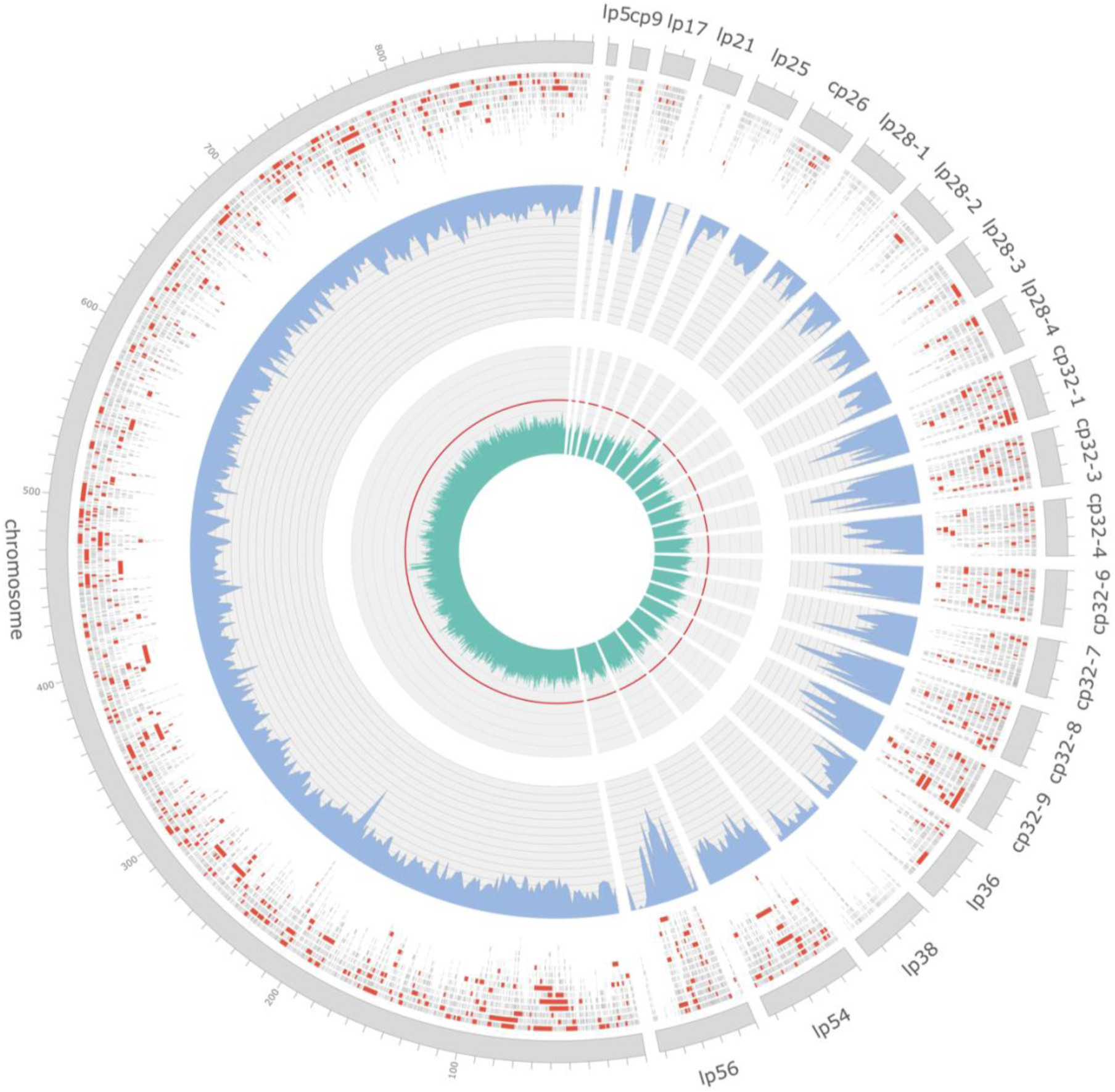
Circular visualization of *Borrelia burgdorferi* B31 reference genome (GCF_000008685) depicting distribution and coverage of raw reads across adaptive sampling experiments. Nanopore reads mapping to the B31 chromosome and associated plasmids are presented along the outer track with individual reads greater than 1 kb in length shown in red. Mean depth of coverage is presented along the middle track in blue, calculated over a 2 kb sliding window, with a total range of 0 – 80× coverage represented across the y-axis. Percentage of GC content across the B31 genome, calculated over a 500 bp sliding window, is depicted along the innermost track in green with the red line denoting the 50% mark.

**Figure 5.**
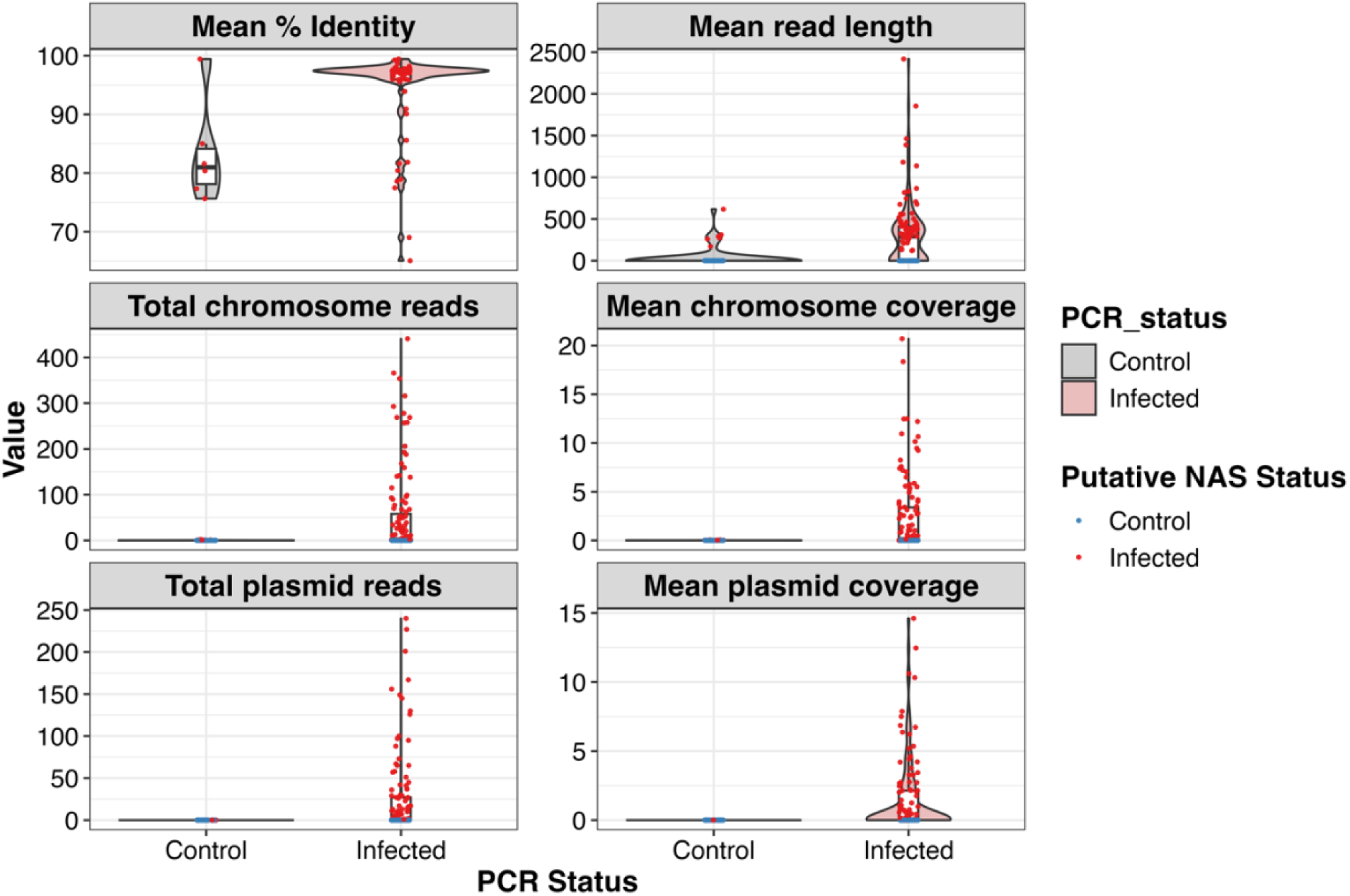
Distribution of read metrics mapping to the *Borrelia burgdorferi* B31 (GCF_000008685) reference genome. Mapped reads are grouped by PCR and putative NAS infection status. Density distributions are shaded by the nPCR status of individual ticks, whereas colored dots represent the putative NAS status of individual tcks.

**Table 4.** Comparison of reads mapping to the *B. burgdorferi* B31 (GCF_000008685) reference genome generated using minimap2, samtools, and seqkit across mapped read filtering criteria. Values represent averages for the cross-tabulated infection status groups. Mapped reads percent identity to the reference genome was calculated using AWK.

|  |  |  |  |  |  |  |  |  |  |  |  |
| --- | --- | --- | --- | --- | --- | --- | --- | --- | --- | --- | --- |
| **Infection status** | **n** | **# of seqs** | **Sum length (bp)** | **Avg length (bp)** | **Max length (bp)** | **N50 (bp)** | **N50 num** | **Q20 (%)** | **Avg quality** | **GC (%)** | **% identity** |
| **B31** |  |  |  |  |  |  |  |  |  |  |  |
| **Raw reads** |  |  |  |  |  |  |  |  |  |  |  |
| NAS+/nPCR+ | 71 | 130.0 | 55,593 | 504.3 | 4679.8 | 1075.3 | 24.7 | 88.1 | 19.3 | 29.2 | 94.6 |
| NAS+/nPCR- | 6 | 1.2 | 369 | 320.6 | 328.0 | 328.0 | 1.0 | 80.2 | 18.8 | 48.7 | 83.2 |
| **> 10 mapped reads** |  |  |  |  |  |  |  |  |  |  |  |
| NAS+/nPCR+ | 56 | 164.4 | 70,279 | 502.5 | 5774.6 | 1208.8 | 31.0 | 87.5 | 18.5 | 27.7 | 96.4 |
| NAS+/nPCR- | 0 | - | - | - | - | - | - | - | - | - |  |
| **Q10 & >300bp** |  |  |  |  |  |  |  |  |  |  |  |
| NAS+/nPCR+ | 65 | 55.1 | 43,132 | 828.2 | 5095.9 | 1465.6 | 12.3 | 88.6 | 19.3 | 28.3 | 95.0 |
| NAS+/nPCR- | 3 | 1.0 | 421 | 421.0 | 421.0 | 421.0 | 1.0 | 78.5 | 17.1 | 53.9 | 79.4 |
| **Q10, >300bp, & >10 mapped reads** |  |  |  |  |  |  |  |  |  |  |  |
| NAS+/nPCR+ | 48 | 73.4 | 57,477 | 847.6 | 6485.7 | 1672.5 | 16.1 | 88.2 | 18.8 | 27.9 | 96.0 |
| NAS+/nPCR- | 0 | - | - | - | - | - | - | - | - | - | - |
| **Q20 & >300bp** |  |  |  |  |  |  |  |  |  |  |  |
| NAS+/nPCR+ | 61 | 32.6 | 24,067 | 761.5 | 3460.3 | 1121.0 | 8.1 | 93.8 | 22.8 | 27.4 | 96.3 |
| NAS+/nPCR- | 1 | 1.0 | 617 | 617.0 | 617.0 | 617.0 | 1.0 | 91.3 | 23.2 | 50.6 | 75.7 |
| **Q20, >300bp, & >10 mapped reads** |  |  |  |  |  |  |  |  |  |  |  |
| NAS+/nPCR+ | 43 | 44.7 | 33,057 | 814.7 | 4484.2 | 1251.5 | 10.8 | 93.7 | 22.5 | 27.5 | 96.6 |
| NAS+/nPCR- | 0 | - | - | - | - | - | - | - | - | - | - |

### Diagnostic Performance Evaluation

Coupling the infection status of individual ticks, as determined using NAS and nPCR, enabled the computation of diagnostic metrics. Contingency tables were constructed across the various filtering thresholds using nPCR as the gold standard, with NAS serving as the diagnostic test result (Supplementary File S2, Table S3). Crude estimation using raw mapped reads revealed high specificity (0.88 [95% CI: 0.76, 0.94]) and PPV (0.92 [95% CI: 0.84, 0.96]) at the expense of sensitivity (0.60 [95% CI: 0.51, 0.68]) and NPV (0.47 [95% CI: 0.37, 0.57]), suggesting an increase in false negatives (Table 5). Filtering for higher quality reads (i.e., Q10 and Q20) led to demonstrable increases in specificity (0.94-1.00) and PPV (0.96-1.00) with concomitant decreases in sensitivity (0.55-0.40) and NPV (0.46-0.41), highlighting the importance of bioinformatic parameterization when ascertaining infection status with sequence data (Figure 6). Filtering for more than 10 mapped reads emphasized this trend, achieving perfect specificity (1.00) and PPV (1.00) across quality filtering thresholds at the cost of diminishing sensitivity and NPV (Figure 6). Across all filtering thresholds, NAS demonstrated consistently high specificity and PPV, averaging 97% (95% CI: 0.93, 1.00) and 98% (95% CI: 0.95, 1.00), respectively, indicating a strong ability to minimize false positives and correctly identify true infections when *B. burgdorferi* reads are detected (Table 5). However, sensitivity varied considerably, averaging 48% (95% CI: 0.41, 0.55) across filtering thresholds, indicating lower detection rates in PCR-positive samples (Table 5). Collectively, increasing the filtering stringency primarily shifted false positives to true negatives, while also shifting some true positives to false negatives (Supplementary File S2, Table S2). The inverse relationship between specificity and sensitivity across filtering thresholds demonstrates the diagnostic trade-off inherent in bioinformatic parameter selection when analyzing NAS sequence data (Figure 7).

**Figure 6.**
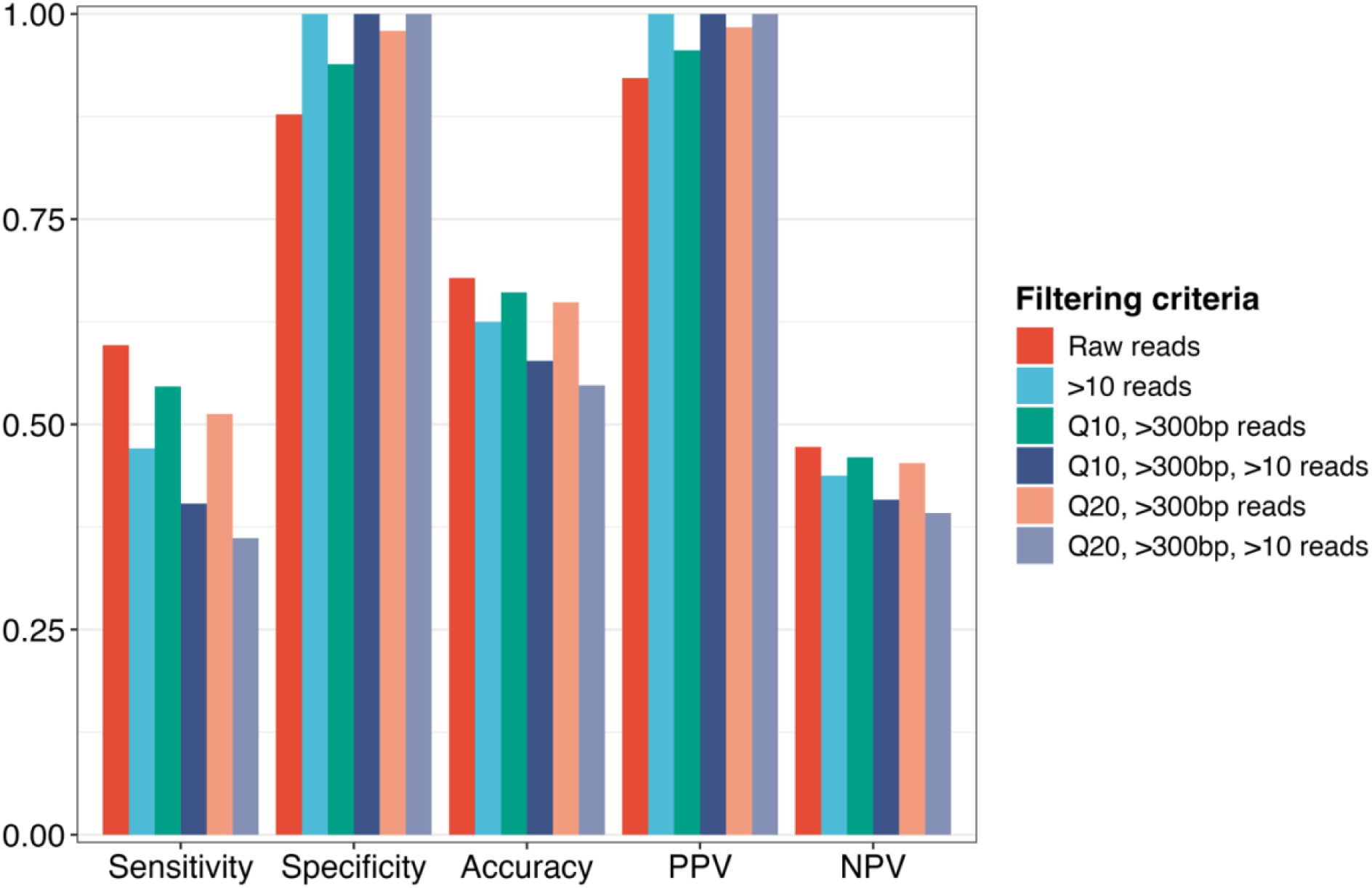
Bar chart comparing the diagnostic metrics of nanopore adaptive sampling for detecting *Borrelia burgdorferi* in wild-caught blacklegged ticks. Colored bars represent diagnostic metrics across filtering criteria. Diagnostic metrics include sensitivity, specificity, accuracy, positive predictive value (PPV), and negative predictive value (NPV).

**Figure 7.**
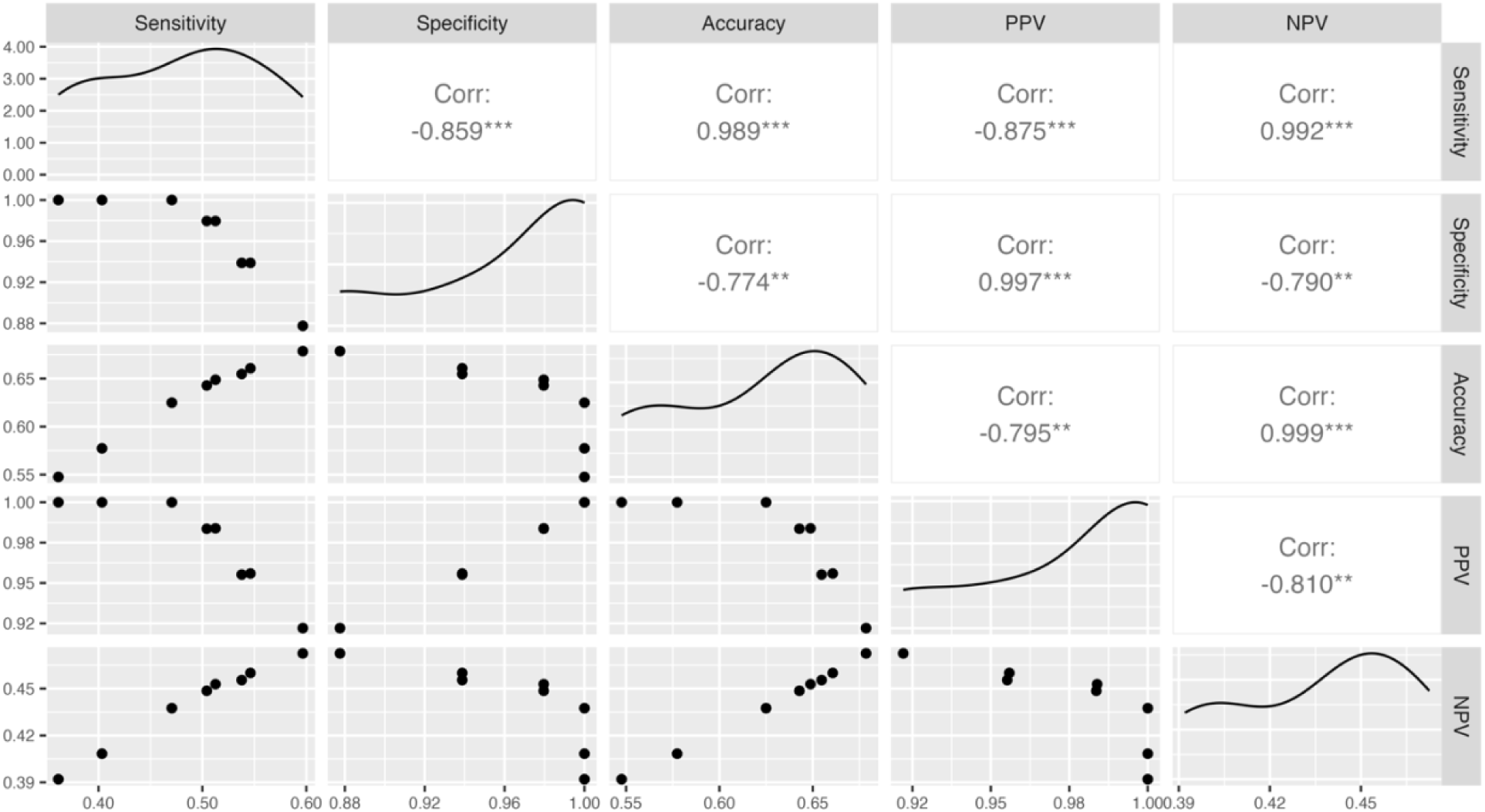
Scatterplot displaying the relationship between diagnostic metrics across the different filtering criteria. The plots running diagonally in the middle of the figure represent the summary correlation curve. The correlation coefficients are displayed for each cross-comparison, with significance evaluated at p=0.05. Asterisks next to correlation coefficients represent the statistical significance of cross- comparisons: *p<0.05; **p<0.01; ***p<0.001.

**Table 5.** Comparison of diagnostic metrics across mapped read filtering criteria with their 95% confidence intervals. 2x2 contingency tables were constructed using nPCR status as the gold standard and putative NAS status as the diagnostic test result. Putative NAS status was determined using reads mapping to the *B. burgdorferi* B31 (GCF_000008685) reference genome.

|  | **Sensitivity** | **Specificity** | **Accuracy** | **Positive Predictive Value (PPV)** | **Negative Predictive Value (NPV)** |
| --- | --- | --- | --- | --- | --- |
| **Mapped to B31** | $TP / TP + FN$ | $TN / FP + TN$ | $TP + TN / n$ | $TP / TP + FP$ | $TN / FN + TN$ |
| Raw reads | 0.60<br>(0.51, 0.68) | 0.88<br>(0.76, 0.94) | 0.68<br>(0.60, 0.74) | 0.92<br>(0.84, 0.96) | 0.47<br>(0.37, 0.57) |
| > 10 mapped reads | 0.47<br>(0.38, 0.56) | 1.00<br>(0.93, 1.00) | 0.63<br>(0.55, 0.69) | 1.00<br>(0.94, 1.00) | 0.44<br>(0.35, 0.53) |
| Q10 & >300 bp reads | 0.55<br>(0.46, 0.63) | 0.94<br>(0.83, 0.98) | 0.66<br>(0.59, 0.73) | 0.96<br>(0.88, 0.98) | 0.46<br>(0.37, 0.56) |
| Q10, 300bp, & >10 mapped reads | 0.40<br>(0.32, 0.49) | 1.00<br>(0.93, 1.00) | 0.58<br>(0.50, 0.65) | 1.00<br>(0.93, 1.00) | 0.41<br>(0.32, 0.50) |
| Q20 & 300bp reads | 0.51<br>(0.42, 0.60) | 0.98<br>(0.89, 1.00) | 0.65<br>(0.57, 0.72) | 0.98<br>(0.91, 1.00) | 0.45<br>(0.36, 0.55) |
| Q20, 300bp, & >10 mapped reads | 0.36<br>(0.28, 0.45) | 1.00<br>(0.93, 1.00) | 0.55<br>(0.47, 0.62) | 1.00<br>(0.92, 1.00) | 0.39<br>(0.31, 0.48) |
| **Average (95% CI)** | 0.48<br>(0.41, 0.55) | 0.97<br>(0.93, 1.00) | 0.62<br>(0.58, 0.66) | 0.98<br>(0.95, 1.00) | 0.44<br>(0.41, 0.46) |

## Discussion

Conventional TBP surveillance relies on targeted PCR assays that limit comprehensive genomic analysis of circulating TBP strains [16]. Although amplification approaches have accelerated the detection of TBPs, nanopore adaptive sampling (NAS) offers unique opportunities for real-time pathogen enrichment and rapid molecular surveillance [23]. However, its diagnostic capabilities, relative to PCR, remain undetermined. Here, we provide the first evaluation of NAS for detecting *B. burgdorferi*, the etiological agent of Lyme disease, in wild-caught blacklegged ticks compared to conventional PCR. We successfully recovered genomic sequences spanning the *B. burgdorferi* genome with distinct differences among PCR-positive and PCR-negative samples, revealing sequence characteristics that distinguish true from false positive detections. This enabled the determination of key diagnostic criteria, such as sensitivity and specificity. Our results demonstrate that adaptive sampling can serve as a rapid, field-deployable tool for TBP surveillance, bridging traditional PCR and genomic approaches.

Our evaluation revealed diagnostic performance capabilities with important implications for TBP surveillance. NAS demonstrated consistently high specificity (97%) and PPV (98%) across filtering thresholds, indicating a strong capability for confirming *B. burgdorferi* presence when detected. However, sensitivity (48%) and NPV (44%) were moderate, suggesting an elevated false negative rate, characterizing approximately half of the PCR-positive ticks not detected by NAS. The trade-off between sensitivity and specificity for molecular surveillance of TBPs ultimately depends on the study objective and surveillance context [27]. For agnostic surveillance aimed at pathogen discovery, higher sensitivity is often preferred, wherein confirmatory methods such as PCR or culturing can help rule out false positives [28]. Conversely, targeted surveillance programs with limited resources may prioritize specificity to maximize the reliability of positive detections [29]. Practically, the burden of Lyme disease necessitates highly sensitive *B. burgdorferi* detection in ticks, and missing infected ticks may skew risk perception and exacerbate disease incidence in emerging or endemic foci. However, over- or underestimation of TBP prevalence due to highly specific or sensitive molecular tools may misdirect key resources necessary to mitigate infected tick encounters among the public. In our study, the moderate sensitivity (48%) suggests NAS may be most suitable for confirmatory surveillance and pathogen characterization. However, optimization of DNA extraction protocols and NAS enrichment will likely enable NAS deployment as an initial field-based tool to generate real-time genomic insights into TBPs. Benchmarking these performance characteristics under different biological contexts and technical parameters is essential for implementing NAS effectively into public health surveillance programs.

The moderate sensitivity observed in our study highlights several technical challenges that may help explain why NAS struggled to detect approximately half of the PCR-positive samples. In the current study, we elected to individually multiplex 24 ticks per sequencing experiment to maximize data generation, minimize per-sample sequencing costs, and mirror practical application in surveillance programs. Pooling individual ticks sacrifices the sequence output of individual ticks as more ticks are pooled, which is heightened if the input gDNA concentrations per tick are not normalized, with downstream consequences on pathogen detection and diagnostic capabilities. Previous research has demonstrated high sensitivity on PCR-confirmed infected ticks when pooling 2 to 4 ticks per flow cell using NAS [23], suggesting that pooling 24 ticks on a single flow cell may have a significant impact on pathogen detection. These pooling constraints are compounded by the use of low molecular weight DNA extraction kits (i.e., spin column-based Qiagen DNEasy preps), producing short DNA fragments consistent with the resulting short read lengths and read N50. Previous research has demonstrated the importance of using high molecular weight DNA extraction kits for long-read sequencing to enhance read lengths and quality [30]. Additional research extending this work should evaluate the trade-off between sample size and the diagnostic capability of NAS while optimizing DNA extraction methods to ensure that input concentrations and qualities are amenable to long-read sequencing.

These sequencing related challenges are exacerbated by the fundamental biology of tick-pathogen systems. Next-generation sequencing generates genomic data on all DNA present in an organism, typically recovering pathogenic DNA at high sensitivity relative to amplification techniques [31]. However, TBPs tend to be found at relatively low copy numbers in unfed ticks, whereas non-pathogenic commensal bacteria tend to be present in high copy numbers, coupled with the overwhelming presence of tick genomic DNA, which creates challenges in recovering sufficient sequence data for confirmatory diagnosis [32,33]. As such, many nPCR-positive ticks likely had very low bacterial loads, making them difficult to detect with our PCR-free NAS approach. Contributing to this difficulty, we chose to compare NAS with nested PCR, which is an extremely sensitive method that could potentially yield more false positives than a conventional, single-reaction PCR. Our observation of the unexpectedly high *B. burgdorferi* prevalence (∼70%) could thus be partly explained by our assay choice. Numerous PCR assays are available to detect *B. burgdorferi* (e.g., 16S, FlaB) that may yield very different diagnostic metrics. Future research should seek to systematically characterize the diagnostic capability of NAS compared against different PCR assays, and ideally, employ qPCR to correlate NAS detection and read statistics with pathogen copy numbers.

Beyond optimizing laboratory protocols, bioinformatic filtering strategies proved helpful in improving diagnostic performance. Filtering sequence data for thresholds of mapped reads completely eliminated false positives, with concomitant increases in specificity and PPV at the expense of sensitivity and NPV, emphasizing the importance of bioinformatic considerations when analyzing NAS enrichment data. Filtering by the number of mapped reads based on true positive distributions is not a deterministic method to eliminate false positives, yet coupling total mapped reads with their underlying sequence characteristics discriminated true and false positives. As the specificity and PPV for raw mapped reads (that included false positives) were 88% and 92%, respectively, it is clear that future research must prioritize strategies to elevate sensitivity and NPV. Sequence characteristics helped to differentiate true and false positive samples, with true positives displaying longer read lengths, greater chromosomal and plasmid coverage, and higher sequence similarity than false positives. Interestingly, mapped read GC percentages for true positives were nearly identical to the *B. burgdorferi* genome, whereas false positives were roughly two-fold greater. False positive ticks contained 1-2 mapped sequences per organism that represent off-target reads from other bacterial taxa, including environmental *Mycobacteria* and *Rickettsia buchneri*, a known commensal endosymbiont of blacklegged ticks. GC content discrimination may be a biological indicator that helps differentiate putative true and false positives, requiring further investigation to clarify this trend. Additional research should prioritize the procurement of NAS enrichment files that counterbalance the detection of numerous TBPs while minimizing off-target read enrichment.

These off-target reads emphasize the challenges in NAS experimental design. NAS presents a novel approach to characterizing pathogenic reads through the selective enrichment of sequences aligning to user-specified referent genomes [20]. However, retaining sequences with lower sequence similarity leads to off-target read enrichment, demonstrated by our finding that false positives represent environmental contamination and commensal endosymbiont infection. In the current study, we included multiple bacterial and protozoal reference genomes in the NAS enrichment file, primarily to capture co-infections within individual ticks, which ultimately may have negatively impacted on-target *B. burgdorferi* mapping rates (0.1%). This broader enrichment approach likely facilitated the capture of high copy number bacteria that dominate tick microbiomes, such as *R. buchneri*, reducing the proportion of reads mapping to low-abundance *B. burgdorferi*. Considerations to maximize data generation by capturing greater genomic diversity while preserving targeted surveillance are imperative when designing NAS experiments.

Developing real-time, unbiased pathogen surveillance systems is critical as invasive tick vectors and emerging TBPs expand throughout North America. NAS provides a platform to characterize the presence, diversity, and distribution of novel tick-borne disease threats by selectively enriching or depleting for target sequences. This enables researchers and public health departments to detect co-infections and putative TBPs simultaneously. Our study provides the first evaluation of NAS’s diagnostic capabilities on *B. burgdorferi-*infected blacklegged ticks in Minnesota. Future research should seek to expand these findings by refining laboratory procedures and streamlining bioinformatic tools for implementation in public health laboratories. These advances will help enhance our capacity to detect emerging tick-borne disease threats and inform public health surveillance strategies.

## Materials and Methods

### Tick collection & DNA extraction

Blacklegged ticks were collected as part of an extensive tick surveillance project conducted across three counties (Anoka, Washington, Winona) in Minnesota. Tick collections involved dragging a 1 m^2^ white cloth along vegetation to capture actively questing ticks. Drag cloths were inspected for ticks every 10 meters; any attached ticks were removed with forceps and placed in 70% EtOH for transportation. Blacklegged ticks were morphologically identified to life stage using the taxonomic keys of Kierans and Clifford [34] and Cooley and Kohls [35]. Total genomic DNA was extracted from adult blacklegged ticks using a Qiagen DNEasy Blood and Tissue Kit (Qiagen, Hilden, Germany) following the manufacturer’s instructions. Ticks were bisected with a sterile 16-gauge needle and incubated in lysis buffer (180 μL Qiagen Buffer ATL and 20 μL Proteinase K) at 56 °C overnight [23]. Genomic DNA was eluted in a final volume of 150 μL and quantified using a NanoDrop One fluorometer (ThermoFisher Scientific, Waltham, MA). Ticks were stored at −20 °C for downstream PCR and sequencing experiments.

### PCR analysis

We leveraged a nested PCR assay (nPCR) to detect *B. burgdorferi sensu stricto* in wild-caught blacklegged ticks [36]. Nested PCR assays were performed in two steps: the first used the extracted tick DNA as a template containing outer primers that targeted a longer amplicon. The second step leveraged the amplicons generated in step one as template DNA, with internal primers targeting a smaller region of the larger amplicon. Our nPCR assay targeted the outer surface protein A (OspA) gene, which encodes a major surface antigen of *B. burgdorferi*, using primers reported in [36]. Detailed information on PCR primers is provided in Table 1. nPCR reactions were performed in 25 μL solutions comprised of 2 μL of template DNA, 12.5 μL of Taq 2X master mix, 0.5 μl each of 10 μM forward and reverse primers, and 9.5 μL of nuclease-free water. Reactions were conducted in a Bio-Rad T100 Thermo Cycler (Bio-Rad Laboratories, Hercules, CA). Thermal cycler conditions were programmed with an initial denaturation step at 95 °C for 5 min, followed by 40 cycles of denaturation at 95 °C for 15 s, 30 s of annealing at 55 °C (step one) or 58 °C (step two), 45 s of extension at 72 °C except for the 40th cycle which lasted for 5 mins and an indefinite hold at 4 °C. Validated *B. burgdorferi* DNA isolates from previous work in our laboratory were used as positive controls in respective nPCR runs [37]. PCR products were run on an electrophoretic gel and visualized using a 150 bp molecular ladder on a 2% agarose gel. Nested PCR procedures were run in duplicate to validate infectivity.

### Nanopore adaptive sampling

#### Sample selection

Following PCR analysis, we randomly selected total gDNA extracts from 70% *B. burgdorferi-*positive and 30% *B. burgdorferi*-negative ticks for nanopore sequencing experiments. Briefly, positive and negative samples were separated into two datasets, and a random subsample was drawn from each. An infection prevalence of 70% and a likely sensitivity of 0.80 (null hypothesis) to 0.90 (alternative hypothesis) requires a total of 153 samples (power: 0.819; p-value: 0.040). Each library preparation kit has the capability to multiplex 24 samples and to ensure a sufficient initial library concentration (> 1 ug), we increased the sample size to 168 total gDNA extracts for sequencing. Each library contained the same proportion (i.e., 70:30) of nPCR-positive and nPCR-negative samples, respectively. To ensure blinding, ticks from negative and positive nPCR-infected classes were selected for a specific run using a random number generator to determine their inclusion and relative barcode order. Samples from negative and positive infected classes were prioritized according to their DNA concentration and quality to maximize the input library concentration for downstream sequencing. All subsequent analyses using sequence data were performed blinded to infection status. ONT native barcoding recommends that each barcode have an initial DNA concentration of 400 ng, and gDNA extracts that did not meet this threshold were subject to vacuum centrifuge concentration using a Vacufuge Plus (Eppendorf). Concentrated gDNA extracts were quantified using a NanoDrop One fluorometer before library preparation. DNA inputs were not normalized before library preparation.

#### Library preparation

Seven individual libraries, each consisting of 24 ticks, were constructed using the ONT Native Barcoding Kit (SQK-NBD114) following the manufacturer’s instructions for use with a MinION flow cell (R10.4.1). All libraries were prepared identically. DNA ends were initially repaired and prepared for barcode and adapter ligation using the NEBNext FFPE DNA Repair Mix and NEBNext Ultra II End Repair/dA-tailing Module (New England Biolabs Inc., Ipswich, MA). Each sample was incubated at 20 °C and 65 °C for 5 minutes, respectively, before purification and concentration using AMPure XP magnetic beads (Beckman Coulter, A63881). A 1:1X ratio of magnetic beads was used at each bead cleanup step. The samples were then cleaned with 80% EtOH and eluted in 10 μL of nuclease-free water. ONT molecular barcodes (Oxford Nanopore Technologies) were added to the eluate, enabling pooling, using a Blunt/TA Ligase Master Mix (New England Biolabs Inc.), and incubated for 20 minutes at room temperature. Following incubation, 4 μL of EDTA (Oxford Nanopore Technologies), which acts as a chelating agent to inhibit exonucleases from degrading the DNA, was added to each sample, and all samples were pooled into a single tube. The pooled, barcoded library was then purified and concentrated using AMPure XP magnetic beads, incubated at room temperature, cleaned using 80% EtOH, and eluted in 35 μL of nuclease-free water. Native adapters (ONT) were then added to the pooled barcoded library, which serve to anchor DNA fragments to individual nanopores on the flow cell for sequencing, using the NEBNext Quick Ligation Reaction Buffer and Quick T4 DNA ligase (New England Biolabs Inc.), and incubated for 20 minutes at room temperature. The library was then cleaned and purified using AMPure XP magnetic beads, incubated at room temperature, and cleaned using the ONT short fragment buffer. The final pooled library was eluted in 15 μL of Elution Buffer (ONT). At each elution step, the libraries were quantified using a Qubit 4 Fluorometer with the dsDNA HS Assay Kit (Thermo Fischer Scientific, Q32854).

#### Sequencing and basecalling

Each barcoded library (n=7) was sequenced using individual MinION Flow Cells (R10.4.1) on a MinION Mk1B instrument (ONT). Sequencing experiments were performed on a desktop computer using Pop!_OS with the following specifications: AMD Ryzen 9 7900X 12-core processor (Advanced Micro Devices, Inc., Santa Clara, United States), NVIDIA GeForce RTX 4090 (Nvidia, Santa Clara, United States), 64 GB RAM, 4 TB SSD, and an 8 TB internal hard drive. Sequence experiments were parameterized and monitored using MinKNOW (ONT). Each flow cell was checked to ensure that an adequate (i.e., >800) number of pores were available for sequencing before loading each of the final libraries. Flow cells were primed with a mixture of ONT Flow Cell Flush and Flow Cell Tether (ONT) that was gently loaded into the priming port, avoiding any introduction of air bubbles. Each library was prepared with ONT Sequencing Buffer and Library Beads (ONT) and then loaded onto the flow cell’s SpotOn sample port. After loading, sequencing parameters were specified using MinKNOW, specifically enabling the enrichment of specific sequences as they pass through individual nanopores, termed nanopore adaptive sampling (NAS). NAS allows users to selectively enrich or deplete for target sequences, including whole genomes. As a DNA strand passes through the nanopore, the electrical signal is disrupted, producing a distinct change that represents the passing nucleotide sequence, which is basecalled in real-time using Dorado v0.8.1 (ONT). Nanopore adaptive sampling takes the first few hundred bases of the passing nucleotide sequence and maps it to the user-specified reference file. When enrichment is selected, if the passing nucleotide sequence is 70% similar to any sequence in the reference file, the sequencing of that DNA fragment will continue to completion. However, if the sequenced fragment is <70% similar, it is ejected from the nanopore, and the pore is freed to sequence a new strand. Here, each sequencing experiment was performed with an enrichment file containing known DNA-based TBPs vectored by the blacklegged tick: *Borrelia burgdorferi sensu stricto* (GCF_000008685), *Borrelia miyamotoi* (GCF_019668505), *Ehrlichia muris euclairensis* (GCF_000508225), *Babesia microti* (GCF_000691945), and *Anaplasma phagocytophilum* (GCF_000439775). Each reference genome FASTA file was downloaded from NCBI, concatenated using the command line into a penultimate reference file, and indexed using minimap2 [21]. The indexed pathogen reference file was selected for enrichment and alignment in MinKNOW before starting each sequencing experiment. Additional sequencing parameters were specified in MinKNOW before sequencing, including read quality (Q>8), minimum read length (>250 bp), the location to write output files, and library preparation kits used to construct libraries. Each flow cell was sequenced to exhaustion (e.g., 72 hours) without washing and reloading individual libraries.

### Sequence data analysis

#### Basecalling and pre-processing

Raw data from each sequencing experiment in POD5 file format were rebasecalled, adapter trimmed, and demultiplexed using Dorado with the “super accuracy” model dna_r10.4.1_e8.2_400bps_supv4.2.0. Passed FASTQ files, which included reads with a quality score greater than 8 and a minimum read length of 250 bp, were extracted and utilized for downstream analysis. Each barcode (n=24 in each of 7 experiments) contained multiple zipped FASTQ files for every sequenced DNA fragment that were concatenated using the command line into single zipped FASTQ files comprising all reads for a given barcode.

#### Bioinformatics

Raw FASTQ files were retained and further filtered for quality and minimum read length to evaluate the influence of read filtering parameters on diagnostic measurements, as outlined in diagnostic metric computation section below. Summary statistics on raw and filtered FASTQ files were generated using seqkit [38], Nanoq [39], and NanoComp [40]. These FASTQ files were iteratively mapped to the *B. burgdorferi* B31 reference genome (GCF_000008685) using custom scripts with minimap2. Mapped read SAM files were parsed to retain only mapped reads, converted to BAM format, and BAM files were sorted and indexed using custom scripts with samtools [41]. Mapped read depth was evaluated using the sorted and indexed BAM files with custom scripts using samtools and output into a collated tab-delimited text file for export. Mapped read coordinates were ascertained by converting BAM files to BED format using custom scripts with bedtools [42]. Mapped read identity was evaluated using custom scripts written in AWK (https://www.gnu.org/software/gawk/manual/gawk.html#Manual-History). Mapped read BAM files were converted to FASTQ format and evaluated for summary statistics using custom scripts with samtools and seqkit. Reads mapping to the B31 reference genomes (GCF_000008685) linear chromosome and extrachromosomal plasmids were partitioned and visualized with Circos [43]. This process was subsequently performed using two other *B. burgdorferi* reference genomes, B31 (GCF_040790805) and Bol26 (GCF_000181575), to determine the influence of the mapping reference on results. These results are available in Supplementary File S2. Attempts to assemble mapped reads using mapped FASTQ files were performed with Flye [44], SPAdes [45], and canu [46], yet read coverage was not sufficient to generate informative contigs.

#### Diagnostic metric computation

To understand the diagnostic capacity of nanopore sequencing, and more pointedly NAS, we determined the putative infection status of individual organisms, blinded to their PCR infection status. Using the raw and filtered mapped reads from the bioinformatics section described above, putative NAS positivity and negativity was ascertained through the number of reads mapping to the *B. burgdorferi* reference genomes across the various filtering thresholds. For example, using raw reads, any number of reads mapping to the *B. burgdorferi* genome were deemed positive (baseline). The filtering thresholds then enabled cross-comparison to determine the optimal post-processing steps. Evaluating *B. burgdorferi* infectivity in blacklegged ticks with nPCR was treated as the gold standard, or disease status, whereas putative NAS infectivity served as the diagnostic test result, or experimental group. This enabled the estimation of diagnostic metrics for NAS detection of *B. burgdorferi* from naturally collected blacklegged ticks in Minnesota. Under this framework, we were able to construct contingency tables ascertaining diagnostic metrics, using the following convention: ticks that were positive from both nPCR and NAS were treated as true positives (TP); ticks that were positive from nPCR and negative from NAS were treated as false negatives (FN); ticks that were negative from nPCR and positive from NAS were treated as false positives (FP); and ticks that were negative from both nPCR and NAS were treated as true negatives (TN). From this, we were able to calculate the sensitivity (TP / [TP + FN]), specificity (TN / [FP + TN]), accuracy ([TP + TN] / n), PPV (TP / [TP + FP]), and negative predictive value (NPV) (TN / [TN + FN]). Diagnostic metrics, including sensitivity, specificity, accuracy, PPV, and NPV, were computed using custom scripts written in R v4.5.0 (R Core Team, 2022). These values were compared across filtering criteria to assess the influence of user-specified thresholds for detecting *B. burgdorferi*.

#### R analysis

Summary statistics for each barcode’s raw FASTQ, filtered FASTQ, and mapped FASTQ were exported in tab-delimited text files to R for collation, organization, and visualization. Summary statistic files were reformatted using dplyr [47] to compare NAS-enriched read statistics across PCR status. Mapped read summary statistics were reformatted using dplyr to compare across levels of putative NAS infection status and PCR status for the various filtering criteria described in the bioinformatics section. The distribution of read mapping metrics by putative NAS infection status and PCR status was evaluated using tidyr [48] and ggplot2 [49]. Diagnostic metric comparisons across filtering criteria were visualized using ggplot2 and ggsci [50]. The relationship between diagnostic metrics for the various filtering criteria was assessed using a correlation plot generated with GGally [51] and scales [52].

All command-line and R analysis code is available in Supplementary File S3.

## Acknowledgments

The authors thank the Washington County Public Health and Environment staff for collecting ticks. We are incredibly grateful to Dr. Christopher Faulk, Dr. Benjamin Cull, and Benedict Khoo for their feedback and suggestions.

## Supporting information

**Supplementary File S1.** NanoPlot figures for each sequencing experiment.

**Supplementary File S2.** Summary statistics, 2x2 contingency tables, and diagnostic metrics for reads mapped to the Bol26 *B. burgdorferi* reference genome.

**Supplementary File S3.** R Markdown HTML file containing all bioinformatic and R analytical pipelines.

## Funding

This research was supported by a grant from the NIH to J.D.O and P.A.L (R01AI155472).

## Conflicts of interest

The authors declare that they have no conflicts of interest.

## Author contributions

J.C.: conceptualization, experimental design, sample collection, sample processing, data analysis and curation, initial write-up, revision; E.J.K.: conceptualization, data analysis and curation, revision; L.E.F.: data analysis and curation, revision; P.A.L.: conceptualization, funding, revision; J.D.O.: conceptualization, experimental design, funding, revision.

## Notes

### Competing Interest Statement

The authors have declared no competing interest.

